# The roles of NADPH oxidases during adult zebrafish fin regeneration

**DOI:** 10.1101/2021.07.13.452203

**Authors:** Kunal Chopra, Milda Folkmanaitė, Liam Stockdale, Vishali Shathish, Shoko Ishibashi, Rachel Bergin, Jorge Amich, Enrique Amaya

## Abstract

Sustained elevated levels of reactive oxygen species (ROS) have been shown to be essential for whole body, appendage and organ regeneration in various organisms, including planarians, *Hydra*, zebrafish, axolotl, *Xenopus*, geckos and mice. In the majority of cases these roles have been shown via the use of NADPH oxidase pharmacological inhibitors, which generally target all NAPDH oxidases (NOXes). To identify the specific NOX or NOXes essential for ROS production during adult fin regeneration in zebrafish, we generated *nox* mutants for *duox, nox5* and *cyba* (a key subunit of NOXes 1-4). We also crossed these mutant lines to a transgenic line ubiquitously expressing *HyPer*, which permits the measurement of ROS levels in adult zebrafish fins. Using this approach, we found that homozygous *duox* mutants have significantly attenuated ROS levels following fin amputation, and this correlated with a significantly diminished rate of fin regeneration. While the other *nox* homozygous mutants (*nox5* and *cyba*) showed less of an effect on ROS levels or adult fin regeneration, *duox/cyba* double mutants showed a more diminished rate of fin regeneration than *duox* mutants alone, suggesting that Nox1-4 do play a role during regeneration, but one that is secondary to that of Duox. This work also serendipitously found that ROS levels in amputated adult zebrafish fins oscillate during the day with a circadian rhythm.

## INTRODUCTION

A major aim in the field of regenerative biology is to uncover the mechanisms employed by various species in the animal kingdom, enabling them to regenerate organs and appendages, and in some cases, entire bodies. Deciphering such mechanisms, especially those that are evolutionarily conserved across phyla, will provide insight into why some animals have greater regenerative potential than others. Importantly, uncovering those highly conserved mechanisms may lead to novel therapies that could awaken a better regenerative response, clinically, in the sister field of regenerative medicine.

Over the past decade, it has become increasingly apparent that elevated reactive oxygen species (ROS) levels play an essential role during whole body, appendage, organ and tissue regeneration across the plant and animal kingdoms, including tomato, *Hydra*, planarians, *Drosophila*, zebrafish, axolotl, *Xenopus*, reptiles and mammals (Baddar et al., 2019; Bai et al., 2015; Gauron et al., 2016; Khan et al., 2017; Larriba et al., 2021; Love et al., 2013; Pirotte et al., 2015; Santabárbara-Ruiz et al., 2015; Wenger et al., 2014; Zhang et al., 2016). A key outstanding question is what are the mechanisms that control ROS production during regeneration. Findings from several model organisms have suggested a critical role for NADPH oxidases (NOXes) in the production of ROS following injury (Gauron et al., 2013; Love et al., 2013). The NOXes comprise a family of transmembrane, sometimes multicomponent enzymes whose primary physiological function is the transfer of an electron across biological membranes onto molecular oxygen to produce superoxide. The superoxide ion, which is a ROS, can then be dismutated to other more stable and readily diffusible ROS forms, such as hydrogen peroxide (H_2_O_2_) (Bedard and Krause, 2007). Most previous work establishing a critical role for NOXes during ROS production and regeneration has been based on the use of pharmacological inhibitors, such as diphenylene iodonium (DPI), apocynin and VAS2870, which lack specificity and selectivity (Freyhaus et al., 2006; Reis et al., 2020). Therefore, to identify the specific NOXes responsible for ROS generation during adult appendage regeneration, we decided to take a genetic approach using zebrafish. All fins of the adult zebrafish are able to regenerate (Nachtrab et al., 2011). We focused on the caudal fin, which is easily accessible and has unlimited regeneration potential (Azevedo et al., 2011). Here we report the generation and characterisation of several mutant alleles of zebrafish *cyba, nox5* and *duox*. Using homozygous mutants for each of these genes, as well as *cyba/duox* double mutants, we investigated the effect of each mutant on ROS production and adult fin regeneration.

## RESULTS

### Molecular characterisation of *nox* mutant alleles

The primary aim of this work was to identify the molecular mechanisms responsible for sustained ROS production during zebrafish caudal fin regeneration. Previous work provides evidence that NOXes play an essential role in ROS production and subsequent caudal fin regeneration, but that work was based on the use of NADPH oxidase inhibitors (Gauron et al., 2013), which inhibit all NOXes (Reis et al., 2020). Thus, we chose a genetic approach as a more specific strategy to pinpoint the NOXes involved in ROS production and subsequently, caudal fin regeneration. Therefore, we sought to obtain or generate mutant alleles for a number of NOXes or their essential subunits (Fig. 1A). One such critical subunit, P22^phox^, is encoded by *CYBA* gene. P22^phox^ plays an essential role in the maturation and structural integrity of NOXes 1-4 (Panday et al., 2015). Like the mammalian *CYBA* orthologues, the zebrafish *cyba* genomic locus contains 6 exons and 5 introns (Nakano et al., 2008; Stasia, 2016). We also generated a mutant allele for *nox5*, which encodes Nox5, a calcium-regulated single subunit NOX enzyme (Panday et al., 2015). NOX5 is present in humans, but is absent in rodents (Jha et al., 2017). The zebrafish *nox5* has 2 splice variants, both of which are protein coding and feature 16 exons. To try to achieve null mutant alleles for *cyba* and *nox5*, we targeted exon 1 of *cyba*, and two sequences in exon 4 (common to both variants) for *nox5* using CRISPR. F_0_ adults were crossed to WT animals and F_1_ *cyba^+/-^* and *nox5^+/-^* animals were identified via sequencing and subsequently via restriction digest with HindIII and HaeIII, respectively (Fig. 1B-C). For *cyba*, we found a 5bp deletion, leading to a frameshift mutation. For *nox5*, a 4bp indel resulted from the CRISPR-induced mutagenesis, resulting in a frameshift mutation. Incrossing F_1_ heterozygotes led to the establishment of stable homozygous mutant lines, hereafter referred to as *cyba^ex1.5bp-/-^* and *nox5^ex4.4bp-/-^*. For *cyba*, we also obtained a nonsense mutant allele, *cyba sa11798*, from the Zebrafish Mutation Project (Kettleborough et al., 2013). *cyba sa11798* is located in exon 4, leading to a T>A transversion (Fig. 1D) causing a premature stop codon (TAG) after the 87th amino acid. Homozygous mutants for *cyba* and *nox5* are viable, fertile, and display no overt phenotypes. The zebrafish genome encodes a single *duox* gene, instead of the two paralogues (*DUOX1* and *DUOX2*) present in tetrapods (Kawahara et al., 2007). We have previously characterised a *duox* nonsense mutant alleles (*duox sa9892*) (Chopra et al., 2019), which also arose from the Zebrafish Mutation Project (Kettleborough et al., 2013). The *duox sa9892* mutation is located in exon 21, resulting in a C>T transition (Fig. 1E) and a premature stop codon (TAG) after the 944th amino acid. Homozygous mutants for this *duox* allele are viable as adults and display congenital hypothyroidism (Chopra et al., 2019).

**Fig. 1.**
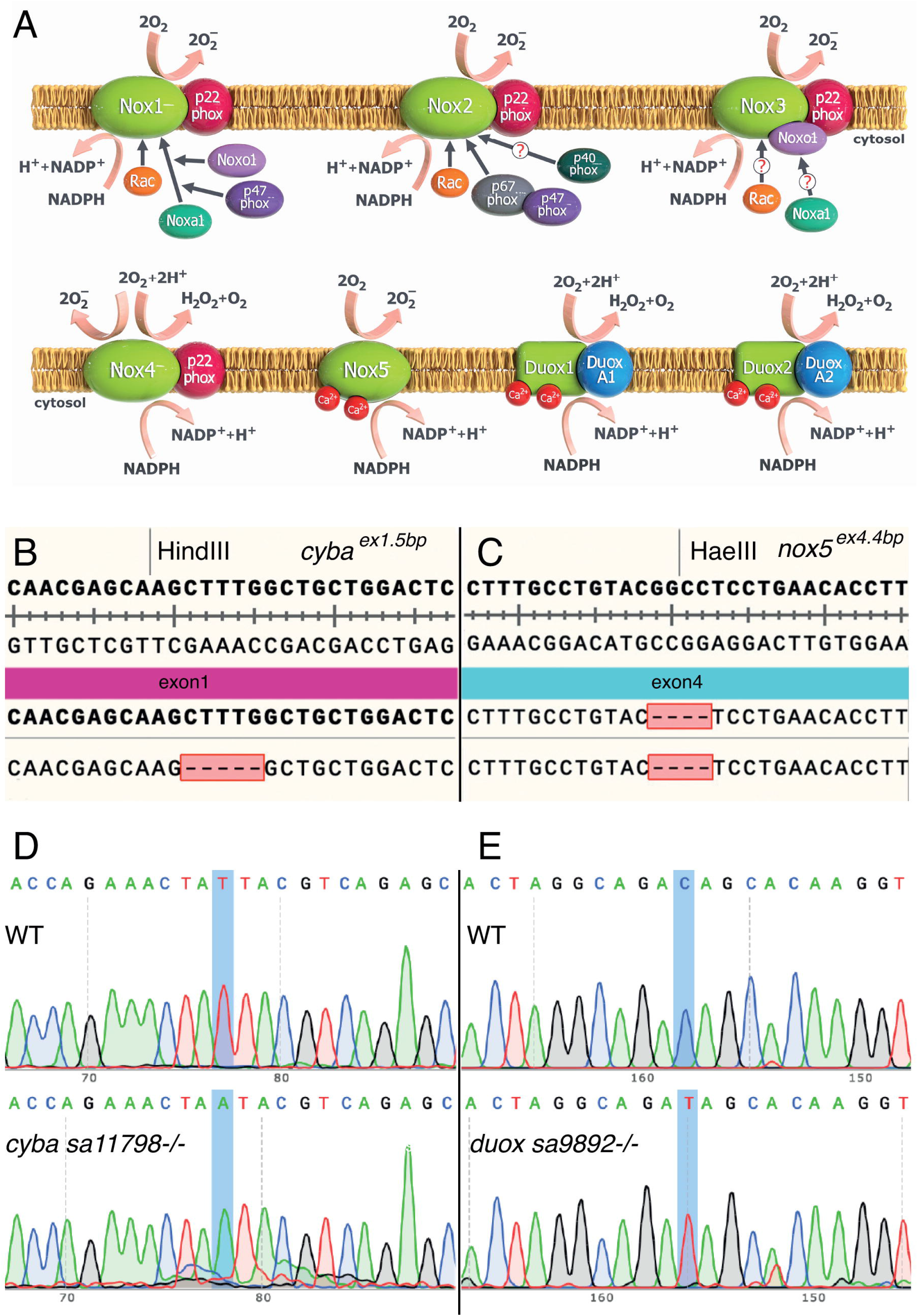
The family of NADPH oxidases. (A) All NOXs are transmembrane proteins that transport oxygen across biological membranes, reducing oxygen to superoxide, which can then undergo dismutation to generate H_2_O_2_. NOXs are multi componential proteins, and p22phox (the product of the *cyba* gene) is a common subunit to (NOX1-4). Zebrafish lack Nox3 and have a single Duox isoform. Characterisation of *cyba sa 11798* (B) and *duox sa9892* (C) via Sanger sequencing shows the single nucleotide changes T>A and C>T, respectively, in contrast to a WT reference sequence. Mutants for *cyba* (D) and *nox5* (E) were generated using CRISPR, resulting in a 5bp deletion and a 4bp indel, respectively. The mutations led to the removal of restriction enzyme sites, providing an easy diagnostic method for screening animals. Panel A courtesy Dr. Kalin Narov, https://kalinnarov.wixsite.com/embryosafari.

### *cyba and nox5* mutations are linked to unique phenotypes

After generating the *cyba, duox*, and *nox5* mutant alleles we then asked whether any of them displayed homozygous mutant phenotypes. We recently showed that *duox* mutant alleles are homozygous viable, but display several phenotypes consistent with congenital hypothyroidism, including shorter body length, external goitres, hyperpigmentation and sterility. These phenotypes could be rescued by T_4_ dosing and could be phenocopied in WT animals via chronic exposure to a goitrogen (Chopra et al., 2019).

For *cyba*, we took clues from its clinical relevance to chronic granulomatous disease (CGD). In human patients CGD is often linked to mutations in the *CYBA* (Stasia, 2016). To this end, CGD patients experience recurrent infections, most frequently brought on by *Aspergillus* spp., *Burkholderia cepacia, Staphylococcus aureus, Serratia marescens, Nocardia* spp. and *Salmonella* (Jones et al., 2008; Marciano et al., 2015; Martire et al., 2008; Winkelstein et al., 2000). This high sensitivity to bacterial and fungal infections arises from inappropriate or insufficient ROS production in innate immune cells, required to kill engulfed pathogens. Although both zebrafish *cyba* mutant alleles are homozygous viable and fertile, we wondered whether they might be more susceptible to fungal infections, and thus exhibit CGD-like phenotypes. Indeed, the *cyba* mutant line we obtained from the EZRC (*cyba sa11798*) had been previously shown to be highly sensitive to fungal infections (Schoen et al., 2019). However, we also wished to confirm that the CRISPR-mediated mutant line we had produced as part of this work (*cyba^ex1.5bp^*) was also sensitive to fungal infections. Given that invasive infections by filamentous fungi are especially important contributors to morbidity and mortality in CGD patients (Beauté et al., 2011; Blumental et al., 2011; Falcone and Holland, 2012), we asked whether *cyba^ex1.5bp-/-^* larvae had increased susceptibility to *Aspergillus fumigatus* infections, relative to WT larvae. To achieve this, we took advantage of the large and easily accessible larval yolk sac to inject *A. fumigatus* conidia in *cyba* mutant and WT larvae. Across two independent experiments (Fig S1 A-B), *cyba^ex1.5bp-/-^* animals (n=25) injected with *A. fumigatus* conidia showed a significantly higher mortality rate (40% in A and 86% in B) at 3dpi, compared to WT larvae (n=17) (17% in A and 27% in B) (Log-rank test; P-value, <0.005) (Fig. S1A-B). Similar results were obtained following injection of *A. fumigatus* conidia into *cyba sa11798^-/-^* mutant larvae (n=8), with mortality rates at 87% in this instance (Fig. S1A). All larvae that developed infections showed similar phenotypes. During the initial stages of infection, the swimming ability of the larvae was significantly affected whereby they could move their pectoral fins but could not achieve productive movement. As the infection advanced, larvae could not remain upright, probably due to effects on the swim bladder (Fig. S1C-D). Terminal infections were characterised by extensive necroses and feebly moving gill arches and heartbeat. Thus, in agreement with the previous study (Schoen et al., 2019), *cyba sa11798^-/-^* animals were very sensitive to *A. fumigatus* infection. Similarly, *cyba^ex1.5bp-/-^* animals were also ultrasensitive to *A. fumigatus* infection. This confirmed that mutations in *cyba* cause high susceptibility to fungal infection in zebrafish larvae, as in human patients with CGD, thus pointing both *cyba* mutant alleles as useful models for human CGD.

Based on the existing literature, we had no clues as to the phenotypes that we might expect in homozygous *nox5* mutant fish. No human disease has yet been associated with *NOX5* mutations, and rodents lack a *Nox5* orthologue. While we found *nox5^ex4.4bp-/-^* fish to be viable and fertile, we serendipitously uncovered an unusual mild phenotype in these animals after treating them with anaesthetics. Intriguingly, most *nox5^ex4.4bp-/-^* animals appeared to have prolonged resistance to the commonly used anaesthetic, tricaine methanesulfonate (MS-222). To characterise this phenotype further, we individually treated *nox5^ex4.4bp-/-^* adults (n=22) and WT adults (n=23) with MS-222 (0.04%) in a beaker. We then measured two responses to the anaesthetic: 1) righting reflex, and 2) cessation of opercular movement. The righting reflex normally serves to resume orientation when the body loses its upright orientation. All WT animals lost the righting reflex in under 30s following immersion (Fig. S2A). After this point, simple tapping of the bench on which the beaker was placed elicited no response from them. In contrast, it took *nox5^ex4.4bp-/-^* animals up to 360s to achieve loss of the righting reflex. Even though they stopped swimming, mutants remained responsive to the tapping, which stimulated jerky, erratic movements. The loss of opercular movement in WT animals took up to 200s, while in the mutants this took as long as 382s (Fig. S2B). Interestingly, most mutants appeared to lose the righting reflex very close to cessation of opercular movement. We further checked the response of both groups to lidocaine hydrochloride and cold water, both of which have been previously tested as anaesthetics in zebrafish (Collymore et al., 2014). While neither group achieved a surgical plane of anaesthesia with 750mg/l of lidocaine, loss of the righting reflex was significantly delayed amongst the *nox5* mutants (Fig. S2A). Meanwhile, cold water induced a rapid surgical plane of anaesthesia on both groups, with no statistically significant difference in their responses (Fig. S2).

Overall, we found that, while all *nox* mutant alleles were homozygous viable into adulthood, each displayed unique phenotypes, suggesting that all mutants were at least hypomorphic, if not null mutant alleles.

### Caudal fin regeneration in zebrafish is not influenced by age

Given that we would be assessing fin regeneration in animals of different genotypes and ages throughout this study, we performed caudal fin amputations in WT fish of different ages to ascertain whether their age might affect the extent and/or rate of fin regeneration. For these analyses, we amputated approximately 50% of the caudal fin in adult animals from 3 months through 22 months of age, and then followed them individually over the four weeks it normally takes for fin regeneration to complete. To gain insight into the rate of regeneration, we imaged the caudal fin of each individual prior to amputation, immediately after amputation and once a week until the end of the fourth week of regeneration (Fig. 2). When plotting the area of the caudal fin as a percentage of its size prior to amputation for each animal at different ages, we found that adult zebrafish younger than six months of age generally overshot the initial size of their fins (i.e. regenerated to over 100% of their initial fin size) (Fig. S3A-C).

**Fig. 2.**
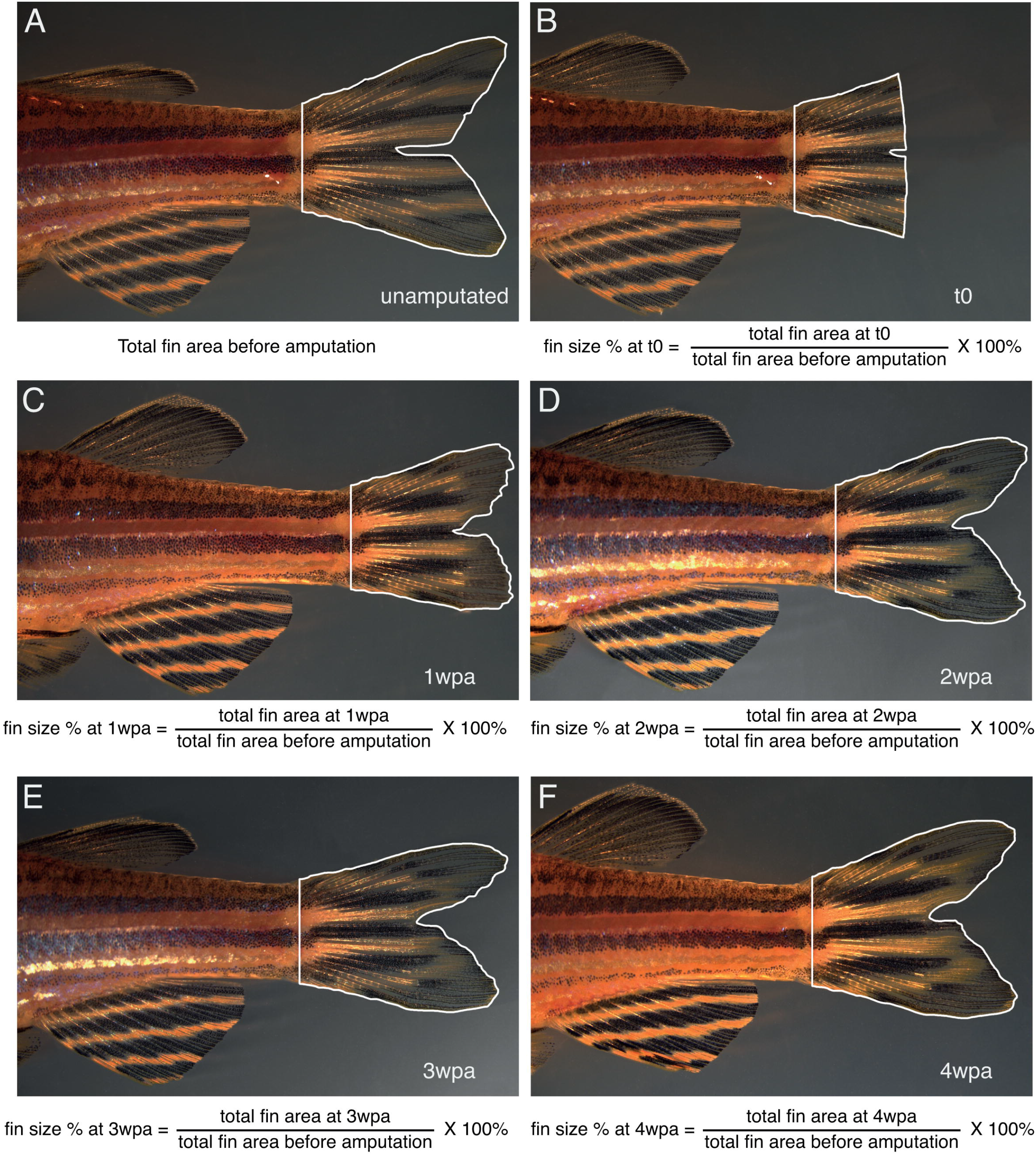
Methodology for quantifying rate of adult fin regeneration. Amputated adult caudal fins in zebrafish achieve complete, scar-free, regeneration by 4 weeks post amputation (wpa). In all cases, the fin was imaged prior to amputation (A), immediately following amputation (B), and then once weekly at 1wpa (C), 2wpa (D), 3wpa (E) and 4wpa (F). The distinctive stripe and pigment patterns on the body and anal fin enabled identification of individual animals. Fins were amputated midway, and the caudal peduncle was used to demarcate the proximal extent. Scale bar=5mm

However, caudal fins in fish six months and older generally only grew back to their original fin size (Fig. S3D-I). From this we can conclude that caudal fin regeneration in young adult fish results from a combination of fin regeneration and growth, while in fish older than 6 months regeneration of the fin is not confounded significantly by fin growth. We thus decided to concentrate the remainder of our tail fin amputation experiments on fish six months or older.

Next, we were interested in addressing whether the rate of fin regeneration is affected by the age of the fish. To answer this question, we plotted the slope of regeneration over the first, second, third and fourth week of regeneration as a function of age (Fig. S4A-E). We found that, overall, caudal fin regeneration is fastest during the first two weeks of regeneration (week 0-1 and week 1-2), and then slows down during the third (week 2-3) and fourth week (week 3-4) (Fig. S4F). Thus, the rate of caudal fin regeneration is biphasic, with similar rates between the first and second weeks, and between the third and fourth weeks, with the latter two statistically slower than the first two weeks (Fig. S4F). Further, we found that animals between six and eight months of age regenerated at the same rate over the four-week time period, while older animals exhibited statistically different regeneration rates relative to animals of other ages during at least one time point (Fig. S4A-E). These results provided us with a good “working age range” of animals, where neither growth nor rates of regeneration were significantly affected by the age of the animals, and thus we concentrated subsequent analyses on animals between six and eight months of age.

### *cyba* and *duox* mutations affect caudal fin regeneration

Having identified regeneration trends in WT animals, we proceeded to investigate the rate of regeneration in various *nox* mutants, namely *cyba^5bp.ex1-/-^* (n=13), *cyba sa11798^-/-^* (n=9), *duox sa9892^-/-^* (n=15), *nox5^4bp.ex4-/-^* (n=6) and (*cyba sa11798^-/-^*)(*duox sa9892^-/-^*) (n=8), along with their WT siblings (n=26) (Fig. 3 and Fig. S5). Interestingly, only *duox sa9892^-/-^* and (*cyba sa11798^-/-^*)(*duox sa9892^-/-^*) animals displayed a significant effect on the rate or extent of fin regeneration. *duox sa9892^-/-^* animals had a significantly slower rate of regeneration relative to WTs and the other single *nox* mutants during the first two weeks following amputation (Fig. 3A). This delayed rate of regeneration was further amplified in the *(cyba sa11798^-/-^)(duox sa9892^-/-^)* double mutants, where the delay in the rate of regeneration continued into the third week (Fig. 3C), although at this point it was only significantly slower than WT and *cyba^5bp.ex1-/-^*. Overall, this reduced rate in the double mutants resulted in stunted regeneration over the entire course of the tail fin regeneration (Fig. 3D; Fig. S5E-F).

**Fig. 3.**
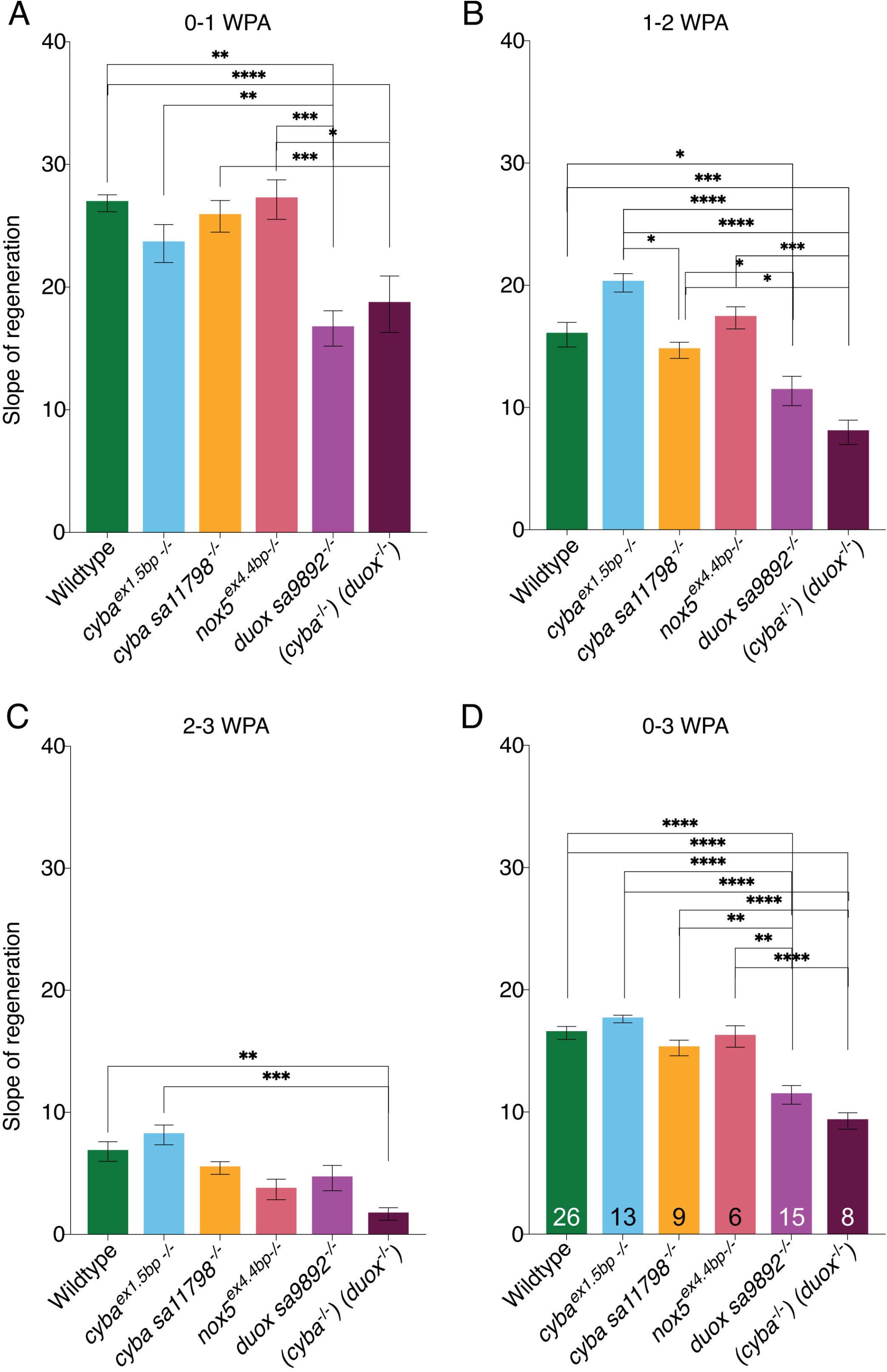
*duox* mutants regenerate the caudal fin slower than WTs. Significant differences in regeneration were observed between WT and *duox* mutants during 0-1wpa (A) and 1-2wpa (B). By 2-3wpa (C) these differences were resolved, with only the double mutant *(cyba sa11798^-/-^)(duox sa9892^-/-^)* continuing to significantly trail behind. (D) An overall view across 0-3wpa highlights how *duox sa9892^-/-^* and *(cyba sa11798^-/-^)(duox sa9892^-/-^)* animals are significantly slower than every other group. Asterisks denote statistically significant differences (Bonferroni’s multiple comparisons test, *P<0.5, **P<0.01, ****P<0.0001).

### The effect of hypothyroidism on the rate of fin regeneration

Recently, we reported *duox* mutant zebrafish as a model of congenital hypothyroidism(Chopra et al., 2019). We thus wondered whether the reduced rate of regeneration among *duox* mutants could be attributed to hypothyroidism. We therefore investigated fin regeneration using another zebrafish model of congenital hypothyroidism, namely the *manet* mutant (McMenamin et al., 2014). The *manet* mutant harbours a nonsense mutation in exon 2 of *tshr*, which encodes the thyroid stimulating hormone (TSH) receptor. We have previously reported phenotypic concordance between *manet* and *duox* mutants (Chopra et al., 2019). Conveniently, the *manet* mutant allele destroys a HpyCH4V restriction site, providing an easy diagnostic method for genotyping. The effect of the *tshr* mutation on body length had not been previously reported, so we measured the body length of WT, *manet^+/-^* and *manet^-/-^* siblings at 3 months of age. This assessment showed a similar growth retardation phenotype we previously reported in *duox* mutants (Chopra et al., 2019) (Fig. S6). We proceeded to amputate the caudal fin in WT, *manet^-/-^* and *duox sa9892^-/-^* fish. In the first wpa, a significantly slower rate of regeneration was observed in *duox sa9892^-/-^*, when compared to WT animals or *manet^-/-^* mutant fish (Fig. 4A). However, during the second and third weeks of regeneration, the *duox sa9892^-/-^* and WT fish had no significant difference in their rates of regeneration (Fig. 4B-C). In contrast, the *manet^-/-^* mutant fish regenerated at the same rate as WT fish during the first week (Fig. 4A), but then their regeneration rate slowed down relative to both WT and *duox sa9892^-/-^* fish during the second and third weeks of regeneration (Fig. 4B-C). In summary, both *duox sa9892^-/-^* and *manet^-/-^* mutants displayed delayed rates of fin regeneration over the three-week period (Fig. 4D). However, the *duox sa9892^-/-^* mutants shows the greatest impact on fin regeneration during the initial phases of regeneration, while hypothyroidism decelerates it only during the later stages, suggesting that the effect *duox* on regeneration is not solely due to their hypothyroidism.

**Fig. 4.**
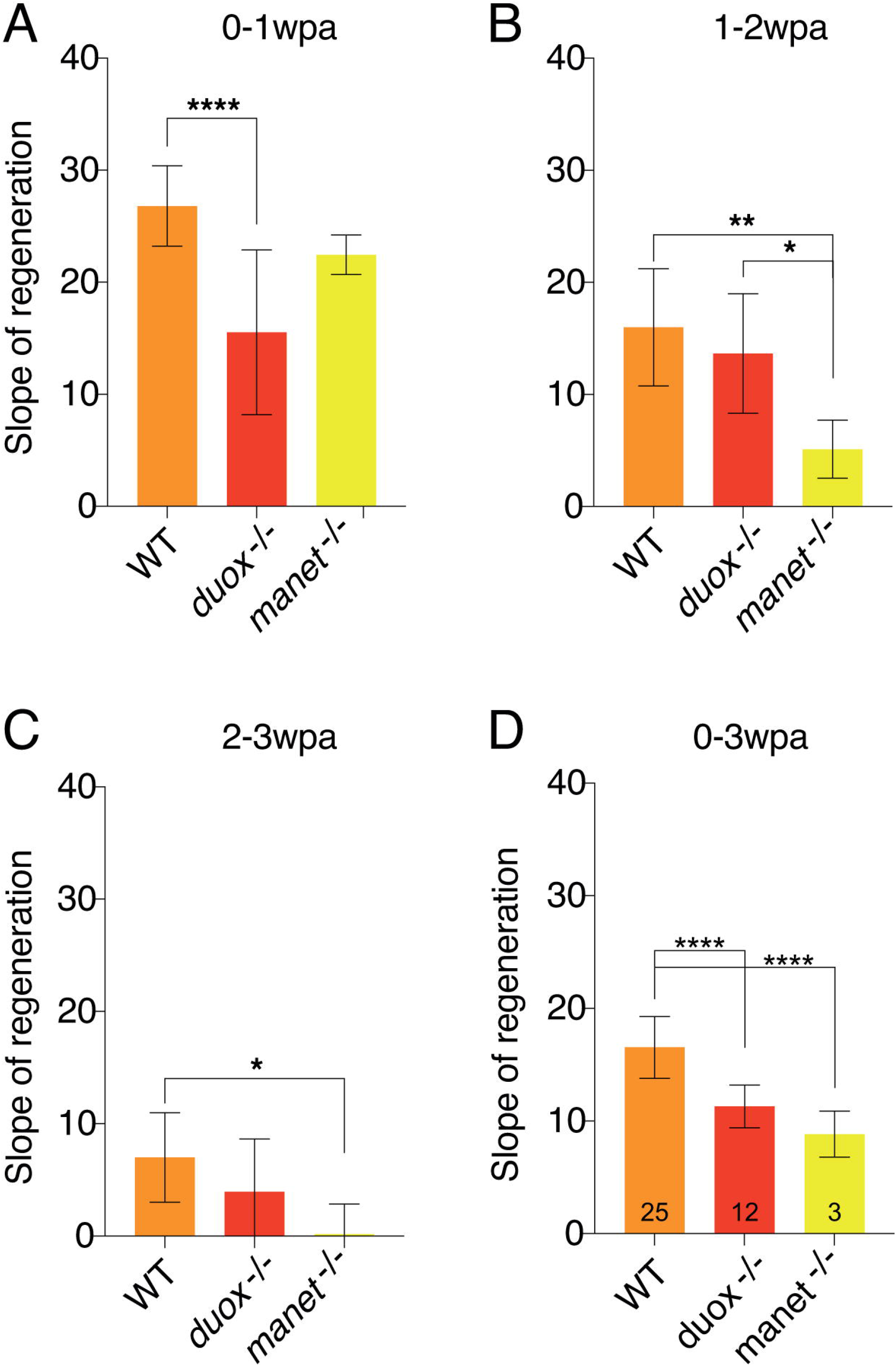
Thyroid hormone deficiency slows down caudal fin regeneration, but only during the late phases of regeneration. (A) The impact of *duox sa9892^-/-^*, but not *manet^-/-^*, on the rate of regeneration is significant during 0-1wpa. By 2wpa (B) the effect of *manet^-/-^* becomes significant, and this effect continues into 2-3wpa (C). (D) An overall view across 0-3wpa indicates *duox sa9892^-/-^* and *manet^-/-^* animals are significantly slower than WTs during regeneration. Asterisks denote statistically significant differences (Bonferroni’s multiple comparisons test, *P<0.5, **P<0.01, ****P<0.0001).

### *ubb:HyPer* is an effective reporter of amputation induced ROS production

Given that the slower rate of regeneration during the first week in the *duox* mutants is not linked to their hypothyroidism, we then asked whether it might be due to a diminished production of ROS in the mutants during this time. To test this hypothesis, we generated a transgenic line that expresses the H_2_O_2_-specific ROS sensor, HyPer (Belousov et al., 2006), under the control of the ubiquitous *ubb* promoter (*ubb:HyPer*). We first measured ROS levels in unamputated caudal fins of individual control *ubb:HyPer* adults (n=7) (Fig. S7A-B), following each animal over the course of four days. Individual animals were identified based on their scale morphology, which is very distinctive under fluorescence. Using a set of repeated time points during the day over four days, we aimed to identify the “baseline” H_2_O_2_ levels in adult fins. While basal H_2_O_2_ levels in uninjured fins were relatively stable throughout the four-day period (Fig. S7B), ROS levels rose significantly above this baseline level following fin amputation (Fig. S7C-D). For consistency, we performed all fin amputations around midday. Observations were recorded at the same time intervals as those used for the unamputated controls, with additional time points at 2, 6 and 10hpa (Fig. S7D). Unlike the unamputated cohort, these were followed for two weeks. By 2hpa (around 3pm), ROS levels showed a significant increase (paired t-test, p=0.0009) above the unamputated level measured at the same time. At 6hpa and 10hpa, the increase in ROS levels was sustained, but thereafter the levels appeared to oscillate each day with the highest levels in the mornings (between 7am and 8am), and the lowest levels in the afternoons (between 3pm and 4pm) (Fig. 5A and Fig. S7D). This rise and fall in ROS levels was recorded over three consecutive days. After three days, the frequency of recordings was reduced to once daily, between 3pm and 4pm. After the sixth day, the frequency of imaging sessions was further reduced to once every two days, between 3PM and 4PM, at which point ROS levels were still higher than the baseline, pre-amputation levels (Fig. S7D). The final recording was made mid-afternoon (between 3pm and 4pm) two weeks after amputation. At this point the ROS levels had finally returned to pre-amputation baseline levels (Fig S7D).

**Fig. 5.**
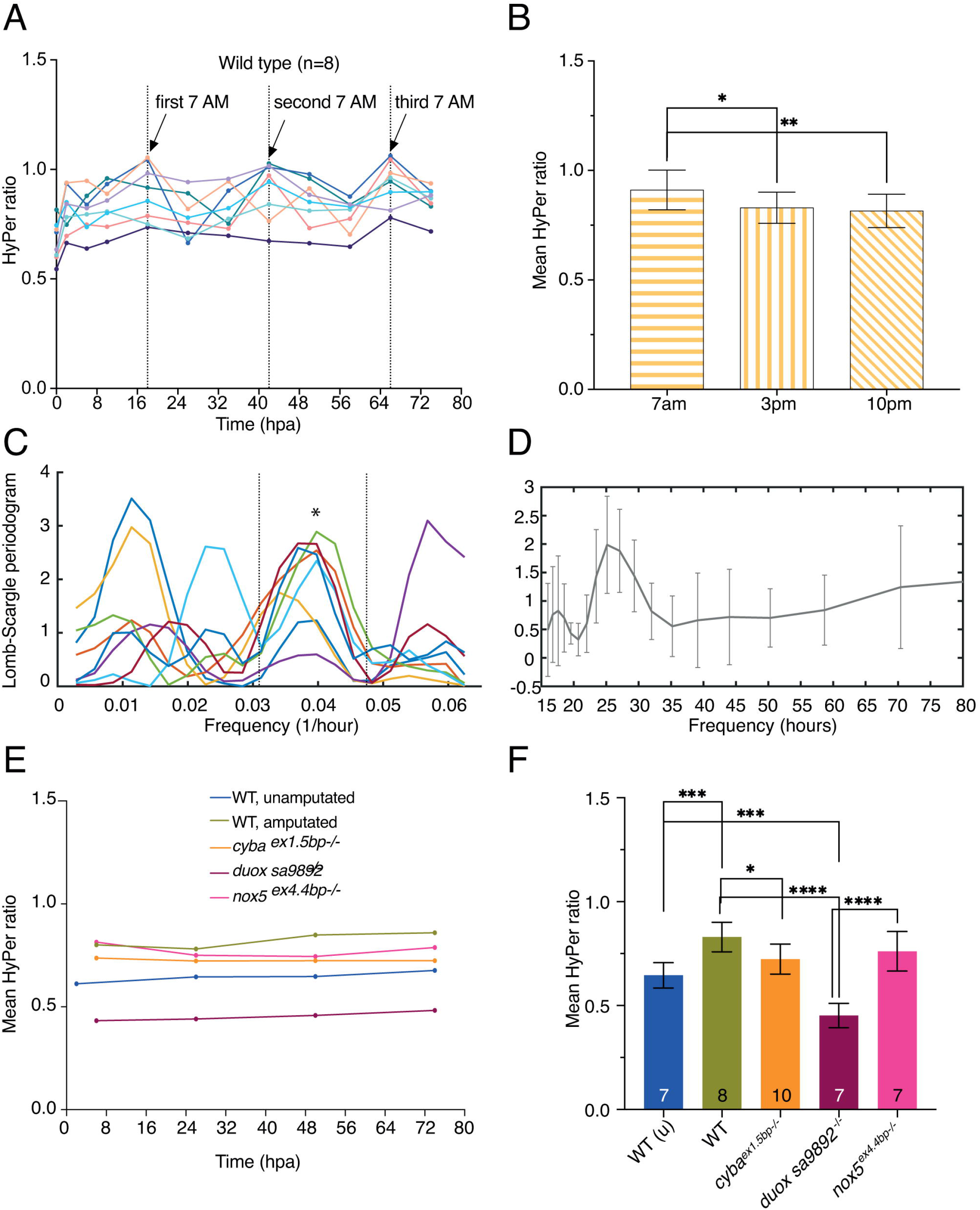
HyPer ratios in WT animals and *nox* mutants. WT animals and *nox* mutants were generated in a *HyPer* transgenic background, which permits *in vivo* assessment of ROS levels. (A) ROS Hyper ratios in individual WT animals (n=8) observed at three time points during the day over three days post amputation reveals an oscillatory pattern of ROS levels. (B) ROS levels over three consecutive days post amputation were significantly higher in the mornings (7am) than in the afternoons (3pm) or nights (10pm). (C) Long-Scargle periodogram reveals that the ROS oscillations follow a daytime-dependent trend, with a periodicity of around 0.4 (* on graph), indicative of a ~24 hour circadian cycle (D). (E) Graph of ROS levels measured at the same time of day (3pm) in WT (unamputated), WT (amputated), *cyba^5bp.ex1-/-^, duox sa9892^-/-^* and *nox5^4bp.ex4-/-^* animals. (F) Graph of mean Hyper ratios measured over three consecutive days at 3pm reveal significantly attenuated post-amputation ROS levels amongst the *duox sa9892^-/-^* mutants relative to the WT amputated fins and the other *nox* mutants. Indeed, ROS levels in the *duox sa9892^-/-^* fins are significantly lower than even those found in unamputated WT fins. ROS levels in the amputated *cyba* mutant fins are significantly lower than those present in WT amputated fins, but not as low as in the *duox sa9892^-/-^* fins. ROS levels in the amputated *nox5^4bp.ex4-/-^* mutant fins are not significantly different to those present in amputated WT fins. (Bonferroni’s multiple comparisons test, *P<0.5, **P<0.01, ****P<0.0001).

We then asked whether the ROS levels measured over consecutive days taken at the same time of the day differed significantly from ROS levels measured at other times of the day. This analysis showed that ROS levels measured at 7am were significantly higher than those measured at either 3pm or 10pm, over the three consecutive days (Fig. 5B). To confirm the veracity of these apparent oscillations we then performed a Fourier transform power spectrum analysis on the changing ROS levels of all the amputated fins over the three days of measurements (Fig. 5C-D). This analysis showed a coordinated high periodicity peak in ROS levels in all fish, at a frequency of 0.04 1/hr (equating to a periodicity of ~24 hours) (Fig. 5C-D). Thus, we found that adult fin amputation led to a sustained increase in ROS levels over the first two weeks of regeneration, and that ROS levels in the amputated fins oscillate with a circadian rhythm.

### Post-amputation ROS production is affected in the *nox* mutants

Having determined the post-amputation sustained increase in ROS levels in wildtype fish (Fig. S7D and Fig. S8A), we next crossed or generated genetically altered lines for all the *nox* mutants in the *ubb:HyPer* background. We then assessed whether post-amputation ROS levels were altered in these mutants. We found that the *cyba^5bp.ex1-/-^* animals (n=10) displayed an increase in ROS levels post-amputation (t-test, p=0.0004) (Fig. S8B). Interestingly, the *cyba* mutants returned to preamputation ROS levels by 1wpa (168hpa) (Fig. S8B), in contrast to the WT animals, where pre-amputation ROS levels were not seen until 2wpa (Fig. S7D). Also, the oscillations in ROS levels during the day exhibited peaks of smaller amplitudes and had a lower H_2_O_2_ increase overall when compared to WT fish. Interestingly, H_2_O_2_ levels in *cyba* mutants were similar to those found in WT fish at 2hpa, but had lower values at later time points.

In contrast, *duox sa9892^-/-^* animals (n=7) (Fig. S8C) showed low ROS levels postamputation throughout the week-long course of observations. The post-amputation levels were significantly lower at all time points taken, and there were no distinctive peaks associated with the 7AM time points. Even more interestingly, ROS levels were lower than those present in WT unamputated controls (Fig. 5D; Fig. S8A and S8C). Finally, the *nox5^ex4.4bp-/-^* animals (n=8) displayed a significant increase in ROS levels at 2hpa (3PM) (t-test, p=0.04), which peaked at 7AM (18hpa) the next morning (t-test, p=0.036), and rapidly ‘baselined’, showing no further significant increase thereafter (Fig. S8D). In order to better compare the HyPer ratios between the different genetic backgrounds we superimposed the ratios obtained for all the genetic strains in Fig. S8E. Given that we previously found significant differences between the HyPer ratios in wild-type animals post-amputation depending on the time of the day, we also compared the HyPer ratios across all the *nox* homozygous mutant strains observed at the same time of day post-amputation (i.e. 3pm) over the first three days post-amputation alongside the ratios found in unamputated tail fins (Fig. 5D). This analysis confirmed that the *duox sa9892^-/-^* mutants had the most profound effect on ROS levels following amputation, where no increase was observed in these animals. Furthermore, it confirmed that the *duox sa9892^-/-^* tail fins have significantly lower ROS levels even in the pre-amputation state. The other *nox* mutants had a much milder effect on post-amputation ROS levels (Fig. 5D).

Overall, these results pinpoint Duox as the primary Nox responsible for ROS production throughout the regeneration period, with the other Noxes also potentially involved in ROS production, but to a much lesser extent. In particular, *cyba* mutants returned to pre-amputation ROS levels after 1wpa, in contrast to wild-type animals, where elevated ROS levels were sustained over the first 2wpa. This may help explain why the *duox* / *cyba* double mutants have a stronger effect on regenerative capacity than the *duox* single mutants during the later phases of tail fin regeneration.

## DISCUSSION

In this work, we set out to uncover the mechanisms of ROS production following zebrafish caudal fin amputation, with particular emphasis on the NADPH oxidase family of enzymes. To address this question, we employed a genetic approach, in contrast to prior studies that employed NADPH oxidase inhibitors (Gauron et al., 2013), which lack specificity. A potential concern in taking a genetic approach was the inherent complication of genetic redundancy often present in teleost fish. Following the divergence of the ray-finned and lobe-finned fishes, teleosts underwent an additional whole genome duplication event in the common ancestor of zebrafish and the other 22,000 ray-finned species (Taylor et al., 2003). Despite this concern we found that the zebrafish genome does not harbour additional duplicated paralogues for *cyba, duox*, and *nox5* when compared to other non-teleost vertebrates. In fact, we and others have found that zebrafish has only a single *duox* gene, as opposed to most other vertebrates, including mice and humans, which have two *duox* paralogues (*DUOX1* and *DUOX2*) (Chopra et al., 2019; Kawahara et al., 2007). Therefore, our work did not suffer from complications arising from excessive genetic redundancy.

Another potential complication could have arisen if homozygous mutants for any of the *nox* genes had been inviable as adults. Fortunately, we found that homozygous mutants for *cyba, duox* and *nox5* are all viable as adults. Although viable, we found that homozygous mutants did display phenotypes, ranging from hypothyroidism and sterility (*duox*), to enhanced susceptibility to fungal infections (*cyba*), and increased resistance to some anaesthetics (*nox5*). Unsurprisingly, the phenotypes associated with *duox* and *cyba* mutations in zebrafish were consistent with mutations in the human orthologues of these genes, with *DUOX2* and *CYBA*, leading to congenital hypothyroidism and chronic granulomatous disease, respectively (Grasberger, 2010; Stasia, 2016). Thus, mutations in the zebrafish orthologues of these two genes provide useful models for these human genetic diseases. In contrast, there are currently no human genetic diseases associated with mutations in *NOX5*, and so we had no prior knowledge of what phenotypes might be associated with mutations in zebrafish *nox5*. In addition, no information can be inferred from mouse knockout studies as rodents lack a *Nox5* gene. Serendipitously, we noticed *nox5^ex4.4bp-/-^* fish were particularly resistant to some anaesthetics, including tricaine and lidocaine. However, we do not currently know the reason behind this. In summary, all the *nox* mutants we used in this work were homozygous viable and each presented with identifiable phenotypes, when appropriately challenged.

The influence of advancing age on regenerative decline is known among adult fish (Itou et al., 2012; Wendler et al., 2015). Given that we would be using adults, we wanted to identify an age range where fin regeneration assays would not be affected by the age of the fish being used. We therefore performed fin regeneration assays, including rate of regeneration, in an array of age groups and found that young adults, from six to eight months of age, showed no significant difference in their rates of fin regeneration. We then set out to assess how each mutation, singly or in combination, affected ROS production following fin amputation, and during fin regeneration. Our results in the *duox^-/-^* mutants were the most striking, resulting in a significantly slower rate of regeneration over the first two weeks following amputation. Furthermore, combinatorial mutants of *duox* and *cyba* displayed an even stronger effect on fin regeneration, extending into the third week following amputation, suggesting that Duox functions in combination with other NOXes during the later stages of adult fin regeneration in zebrafish. Given that the *duox* mutants exhibit hypothyroidism, we also assessed the effect of hypothyroidism alone (i.e. *manet* mutants) on the rate of fin regeneration. While *manet* mutants exhibited slower rates of regeneration, this effect was only apparent during the second week of regeneration. In contrast, the *duox* mutants showed delayed regeneration even during the first week of regeneration and this early delay in regeneration was associated with diminished ROS production in the mutants before and after amputation. These decreased levels of ROS production continued throughout the first week following injury. Previously, genetic approaches have also identified a primary role for the Duox maturation factor in ROS generation during *Drosophila* imaginal wing disc regeneration (Khan et al., 2017), suggesting this role for Duox in regeneration may be a conserved, ancestral function. This is likely due to the fact that Duox is regulated by calcium and a rise in intracellular calcium is a conserved rapid response following injury (Niethammer et al., 2009; Razzell et al., 2013; Soto et al., 2013).

While the focal point of this work was the genetic dissection of the role of NOXes in ROS production and adult fin regeneration, our work uncovered some unexpected findings. One was that ROS levels not only rise after fin amputation, but also the levels oscillate with a circadian rhythm in the days following injury, with the highest levels found in the mornings (around 7am) and the lowest levels in the mid-afternoons (around 3pm). Discovering this was important, as it called for the need to measure ROS levels at the same time daily, for a more accurate assessment of post-amputation ROS levels in the WT versus mutant fish, without being complicated by the circadian-associated oscillatory changes at different times of the day. However, this finding raises several questions. For example, what is responsible for the circadian associated oscillations in injury-induced ROS levels? One possible link might be the well appreciated oscillations in metabolism that are known to be linked to the circadian clock (Panda, 2016). We noted that these oscillations were mostly apparent following injury, which might be linked to the injury induced changes in metabolism that have previously been highlighted in both *Xenopus* tail regeneration and zebrafish heart regeneration (Honkoop et al., 2019; Love et al., 2014). It is thus possible that the oscillating levels of ROS are direct or indirect outputs of the metabolic state in the fin following injury. It is also notable that wound healing in mice and humans is affected by the time of day that the injuries are incurred, or if the circadian clock is disturbed (Brown, 2014; Hoyle et al., 2017; Kowalska et al., 2013; Sandvig et al., 2009; Xue et al., 2017). It will be interesting to investigate whether the time of fin amputation affects wound healing, or the speed, or quality of regeneration in zebrafish, and if so, whether these effects are linked to the dynamics or peak levels of ROS production following injury.

## Materials and Methods

### Zebrafish husbandry

Zebrafish husbandry was undertaken in a re-circulating system maintained at 28.5°C, with a 14hr photoperiod. These conditions are uniform for wild type and all GM strains. Embryos were obtained by marbling tanks, or by isolating pairs in breeding chambers. All animal work was overseen by a Home Office Licence.

### Genomic Extraction

Fin clips from caudal fins of individual adults were added to a mixture of lysis Buffer (10mM Tris-HCL, pH 8.0, 1mM EDTA, 0.3% Tween-20, 0.3% NP40), and proteinase K (20-25mg/ml) (New England Biolabs^®^ Inc.), 1ul/50ul lysis buffer. This was incubated in a thermal cycler programmed to 55°C (2hours), 95°C (10 minutes) and a 12°C hold.

### sgRNA design and production of CRISPR mutants

Single guide RNAs (sgRNAs) were designed for targeting exon 1 of *cyba*, and exon 4 of *nox5*, as previously described (Moreno-mateos et al. 2015). Briefly, the Ensemble ID for each gene when entered online on CRISPRscan generated multiple gRNAs. Exon-targeting gRNAs were then chosen based on rank, location within the first 50% of the ORF, and distance from the initiation codon. A sgRNA template requires a 52nt oligo (sgRNA primer) 5’ TAATACGACTCACTATAGG (N=18) GTTTTAGAGCTAGAA, containing the T7 promoter, the 20nt specific DNA-binding sequence [GG(N=18)] and a constant 15nt tail annealing sequence. This was annealed to an invariant 80nt reverse oligo AAAAGCACCGACTCGGTGCCACTTTTTCAAGTTGATAACGGACTAGCCTTATTTT AACTTGCTATTTCTAGCTCTAAAAC 3’ tail primer, generating a 117bp PCR product. Oligos were obtained from Sigma-Aldrich^®^. The PCR cycler settings for this primer extension were 3 min at 95°C; 30 cycles of 30s at 95°C, 30s at 45°C and 30s at 72°C; and a final step at 72°C for 7 min. PCR products were purified using Qiaquick (Qiagen) columns. Approximately 120–150ng of DNA was used as a template for a T7 *in vitro* transcription reaction. *In vitro* transcribed sgRNAs were treated with DNase and precipitated using sodium acetate and ethanol. Purified sgRNA and Cas9-NLS protein (New England Biolabs^®^ Inc.) were diluted to 300ng/ul.

Equal volumes of Cas9-NLS protein, sgRNA and Phenol Red (Sigma-Aldrich^®^) were mixed to obtain the final injection mix. Injection drop size was adjusted to 1nl using a graticule scale. All embryos were injected at the one-cell stage. F_1_ heterozygous animals were identified via restriction digest, and indels were characterised using Sanger sequencing.

### ENU mutant strains and transgenic strains

ENU mutants for *cyba* (*sa11798*) and *duox* (*sa9892* and *sa13717*) were discovered during the Zebrafish Mutation Project (Kettleborough *et al*., 2013). These fish were sourced from the European Zebrafish Resource Centre (EZRC). *manet* mutants were sourced from the laboratory of David Parichy (McMenamin et al. 2014). *cyba* and *duox* mutants were identified via Sanger sequencing. *manet* mutants were identified via restriction digest. The *Tg(HyPer)* reporter line was generated in the lab using Tol2 transgenesis. The expression of *HyPer* was driven by the ubiquitous promoter *ubb* (Shoko Ishibashi, unpublished).

### Polymerase Chain Reaction

PCRs for were undertaken using ExTaq DNA Polymerase (TaKaRa). Primers used are listed below.

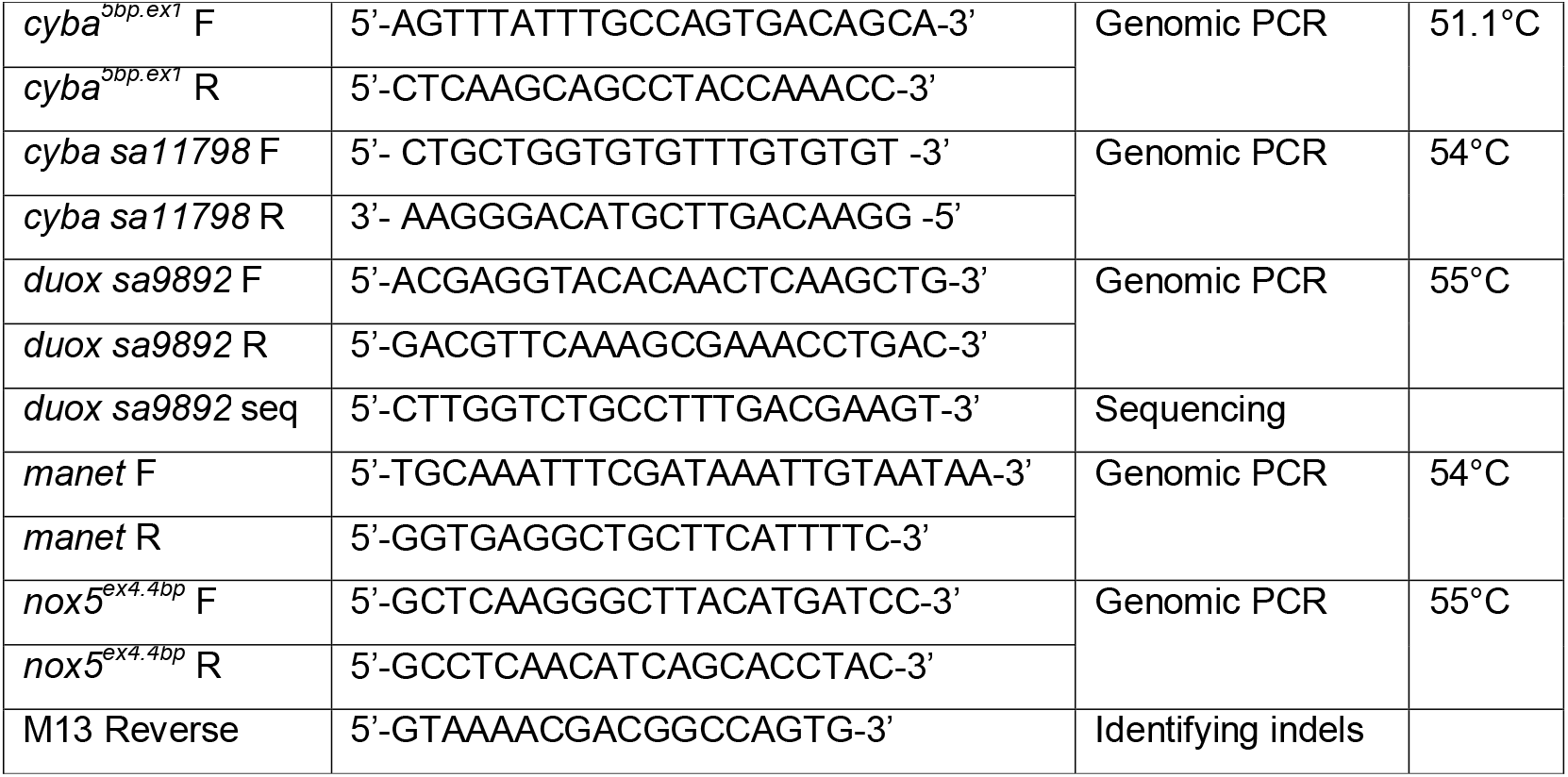

### *Aspergillus fumigatus* spore harvesting and microinjection

An *Aspergillus fumigatus* strain constitutively expressing the Turbo635 fluorescent protein in the A1163 background (Macdonald et al., 2019) was used to infect the larvae. The strain was cultured on *Aspergillus* complete media (ACM) in a 20ml flask and incubated at 37°C for at least 2 days prior to spore harvesting. ACM (1L) was prepared using adenine (0.075g), glucose (10g), yeast extract (1g), bacteriological peptone (2g), casamino acids (1g), vitamin solution (10ml), salt solution (20ml), ammonium tartrate (10ml from 500mM stock), and 1.5% of agar. The pH was adjusted to 6.5 using 10M NaOH. Spores were harvested using a 0.5% NaCl / 0.002% Tween-20 solution (1X). 10ml of the 1X solution was added per flask and flasks were agitated to detach spores. This suspension was filtered through miracloth (Merck Millipore), followed by centrifugation at 4000RPM, for 5minutes. The resultant pellet was resuspended, followed again by centrifugation. Pellet was resuspended in 5ml of the 1X NaCl/Tween. 1/10 and 1/100 dilutions were made from this master solution using and were counted using a New Improved Neubauer haematocytometer (Marinfeld,Germany) on a Nikon Optiphot at a 40x magnification

Freshly harvested spores were used for each experiment. Prior to injection, larvae were anaesthetised in MS-222, and positioned laterally on injection plates made of 3% agarose in E3 (Brothers, Newman, and Wheeler 2011; Knox et al. 2014; E. E. Rosowski et al. 2018). A spore suspension containing 5 conidia/nl was directly injected into the yolk sac of 2dpf larvae, using a microinjection setup consisting of a Picospritzer II (General Valve Corporation) and a Leica MZ6 stereomicroscope. The injected volume was 1nl, unless specified otherwise. Viability of injected larvae was checked twice per day to monitor survival. Dead larvae were plated on Potato Dextrose Agar (PDA). Plates were incubated at 37°C and appearance of hyphal masses was used as confirmation for aspergillosis-led mortality.

### Caudal fin amputation and regeneration

Fish were anaesthetised using 0.4 mg/mL (0.04%) MS-222 (Sigma Aldrich). Animals were imaged prior to amputation, at T0, 1, 2, 3, and 4weeks amputation (wpa), on a Leica M165 FC (Leica Microsystems). Using Photoshop CS5 (Adobe®), images were cropped at the level of the caudal peduncle, and fins were outlined using the brush tool. The “Record Measurements” command was used to obtain the total area of the outlined fin. Regeneration was calculated as a weekly increase, relative to the unamputated state, where the unamputated state was regarded as 100%. This is formulated as:

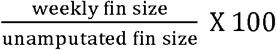

### H_2_O_2_ detection

For visualising H_2_O_2_ levels in the caudal fin, excitation was achieved using BP 430/24 and BP 500/20 filters. Emission was detected with a BP 535/30 filter. Fish were imaged on a high-end widefield microscope (Decon Vision) using a 4X objective. Animals were imaged before and after amputation, over a series of time points. Using ImageJ images were analysed by subtracting background, smoothing, formatting to 32-bit, then dividing to calculate the HyPer ratio. The average H_2_O_2_ over the area captured in the image was then calculated and visualised using Prism 8.1 (GraphPad Software, Inc.).

### Statistical analyses

Aspergillosis-related survival was statistically assessed using the Log Rank Test, in Prism 8.1 (GraphPad Software, Inc.). For fin regeneration, comparisons were made using ANOVA (repeated measures and one-way) or t-test. Linear regressions were performed without interpolation, with a 95% significance level. The strength of the correlation was assessed by the correlation coefficient, adjusted R square. Where applicable, Pearson’s correlation coefficient and Spearman’s rank correlation coefficient was reported. For the unamputated fin, distribution was used for explaining the relationship between age and fin size. Oscillations were tested using power spectrum analysis based on Fourier transform as well as autocorrelation. MATLAB (MathWorks) was used to perform power spectrum analysis and SSPS (IBM Corporation) was used for autocorrelations.

## Acknowledgements

We would like to thank the aquarium staff in the BSF unit for their care and support of the fish. We would also like to thank Kalin Narov, (https://kalinnarov.wixsite.com/embryosafari) for his contribution to Panel A in Figure 1, and Veronica Biga for her help with statistical methods. The Bioimaging Facility microscopes used in this study were purchased with grants from BBSRC, Wellcome and the University of Manchester Strategic Fund. Special thanks go to Peter March, Roger Meadows and Steven Marsden for their help with the microscopy.

## Author Contributions

Conceptualization: E.A., K.C., J.A.; Methodology: K.C., S.I., M.F., L.S., V.S., R.B.; Validation: K.C.; Formal analysis: K.C.; Investigation: K.C., S.I., M.F., L.S., V.S., R.B.; Writing - original draft: K.C.; Writing - review & editing: E.A.; Visualization: E.A.; Supervision: E.A., J.A.; Project administration: E.A.; Funding acquisition: E.A.

---

**Fig. S1.** *cyba* mutants are highly vulnerable to aspergillosis. (A and B) Survival of *cyba^5bp.ex1-/-^* and *cyba sa11798^-/-^* larvae was significantly affected following conidia injection (Log Rank Test, P< 0.005). (C and D) Mutant larvae show systemic infection following *Aspergillus* injection. Conidia injected animals show necroses and the swim bladder (SB) is not distinguishable at 2-3dpi.

**Fig. S2.** *nox5* mutants have a delayed response to some anaesthetics. (A) *nox5^4bp.ex4-/-^* adult animals take significantly longer to respond to agents acting via ion channels, but not to cold water. (B) Following immersion in anaesthesia, but not cold water, opercular movement also continues for longer in *nox5^4bp.ex4-/-^* animals. (Unpaired t test **** P< 0.0001)

**Fig. S3.** Qualitative representation of caudal fin regeneration amongst individual WT adults of various ages is represented using linear regression. Younger animals, especially at 3 and 4 months of age (A and B) exceeded the original unamputated area by 4wpa, while 8 month old and older (E-I) animals regenerate to less than 100% of the original size by 4wpa. This is likely due to younger animals still growing in body size during their period of regeneration. Within each group, individual slopes were not significantly different.

**Fig. S4.** The effect of age on the rate of caudal fin regeneration. No significant different in the rate of regeneration was seen in the first week (0-1) (A) and third week (2-3) (C), but significant differences in various age groups were observed during the second week (1-2) (B) and fourth week (3-4) (D) of regeneration. (E) Taken as a whole, differences in the rates of regeneration were non-significant between most age groups. Majority of the recovery is indicated to occur within the 2wpa, with significant differences recorded between the first 2 weeks and the latter 2 weeks. (Bonferroni’s multiple comparisons test, *P<0.5, **P<0.01, ***P<0.001, ****P<0.0001)

**Fig. S5.** Quantitative representation of caudal fin regeneration amongst individual WT and *nox* mutant adults. Over a three week period, the majority of *duox sa9892^-/-^* (E) and *(cyba sa11798^-/-^)(duox sa9892^-/-^)* fish (F) failed to complete regeneration of the their fins.

**Fig. S6.** External phenotypes are concordant amongst *duox^-/-^* and *manet^-/-^* animals. (A) *duox^+/-^* and *manet^+/-^* zebrafish are similar to WT siblings in terms of body-length, at 3 months of age, while *duox^-/-^* and *manet^-/-^* animals are significantly shorter than all other groups. (B) Quantification of body lengths of WT, *duox^+/-^, duox^-/-^, manet^+/-^* and *manet^-/-^* fish at 3 months reveals a significantly shorter body length in the *duox^-/-^* and *manet^-/-^*, relative to the other genotypes (Bonferroni’s multiple comparisons test, ****P<0.0001).

**Fig. S7.** HyPer ratios in unamputated and amputated WT adult fins over time. (A-B) Hyper ratio is low and remains low over time in unamputated WT fins. (C-D) Hyper ratios rise following amputation and remain higher than baseline levels for up to two weeks following injury.

**Fig. S8.** Average HyPer ratios following amputation in WT and *nox* mutant adult fins over a week (168hpa). *nox* mutants were generated in a *HyPer* transgenic background. Average Hyper ratios in (A) WT, (B) *cyba^5bp.ex1-/-^*, (C) *duox sa9892^-/-^* and (D) *nox5^4bp.ex4-/-^* adult fins. Pink dots correlate with ROS levels at 3pm each day. (E) Average Hyper ratios of all different genotypes shown in the same graph, revealing that *duox sa9892^-/-^* animals have a significantly lower ROS levels than the other genotypes, and also in unamputated WT fins.

